# Patient-derived xenografts undergo murine-specific tumor evolution

**DOI:** 10.1101/167767

**Authors:** Uri Ben-David, Gavin Ha, Yuen-Yi Tseng, Noah F. Greenwald, Coyin Oh, Juliann Shih, James M. McFarland, Bang Wong, Jesse S. Boehm, Rameen Beroukhim, Todd R. Golub

## Abstract

Patient-derived xenografts (PDXs) have become a prominent model for studying human cancer *in vivo*. The underlying assumption is that PDXs faithfully represent the genomic features of primary tumors, retaining their molecular characteristics throughout propagation. However, the genomic stability of PDXs during passaging has not yet been evaluated systematically. Here we monitored the dynamics of copy number alterations (CNAs) in 1,110 PDX samples across 24 cancer types. We found that new CNAs accumulated quickly, such that within four passages an average of 12% of the genome was affected by newly acquired CNAs. Selection for preexisting minor clones was a major contributor to these changes, leading to both gains and losses of CNAs. The rate of CNA acquisition in PDX models was correlated with the extent of both aneuploidy and genetic heterogeneity observed in primary tumors of the same tissue. However, the specific CNAs acquired during PDX passaging differed from those acquired during tumor evolution in patients, suggesting that PDX tumors are subjected to distinct selection pressures compared to those that exist in human hosts. Specifically, several recurrent CNAs observed in primary tumors gradually disappeared in PDXs, indicating that events undergoing positive selection in humans can become dispensable during propagation in mice. Finally, we found that the genomic stability of PDX models also affected their responses to chemotherapy and targeted drugs. Our findings thus highlight the need to couple the timing of PDX molecular characterization to that of drug testing experiments. These results suggest that while PDX models are powerful tools, they should be used with caution.

---

Cancer research relies on interrogating model systems that mirror the biology of human tumors. Cell lines cultured from human tumors have been the workhorse of cancer research for many years, but the marked differences between the *in vitro* cell line environment and the *in vivo* tumor environment raise concerns that cell lines may not be fully representative of human tumors. Recently, there have been increasing efforts to utilize patient-derived xenografts (PDXs) as models to study drug response ^1-4^. These *in vivo* models are assumed to capture the cellular and molecular characteristics of human cancer better than simpler cancer model systems such as established cell lines grown *in vitro* and *in vivo* as xenografts ^1,2^. The value of PDX models for cancer research thus depends on their faithfulness in representing the biological features of primary tumors.

There are reasons to suspect PDX models might deviate from human tumor biology, as they must be serially transplanted for multiple generations in the murine microenvironment. Therefore, it is important to assess whether PDXs retain their genomic and phenotypic characteristics throughout propagation. However, a large-scale systematic analysis of PDX genomic landscapes throughout passaging has not been reported thus far. To date, the genomic stability of PDX models has been primarily evaluated indirectly, leading to the notion that PDXs are highly stable ^3,5,6^. Consistent with this perception, PDX-based studies often involve the analysis of tumors from multiple passages. For example, a recent study of drug sensitivity in PDX models involved the use of tumors varying from passage 4 to passage 10 (ref ^3^).

Hints that PDXs may be more genomically unstable than assumed have begun to emerge, with a recent study showing that the clonal composition of breast cancer PDXs evolves during serial passaging *in vivo* ^7^. Another study recently extended this analysis to additional breast cancer PDXs, showing that while there was overall similarity of PDX models to their tumors of origin, the clonal composition of the tumors could change dramatically throughout PDX derivation and propagation ^8^. Importantly, both studies presented a deep characterization of PDX models from a total of 83 models of a single tissue type (breast), with no systematic assessment of the rate, prevalence and recurrence patterns of genomic changes during *in vivo* passaging of PDXs. Additionally, whether the observed clonal dynamics have any functional importance remains an open question.

Somatic copy number alterations (CNAs) are detectable in the vast majority of cancers ^9,10^, and therefore represent a powerful strategy to track the clonal evolution of tumors. Moreover, CNAs are often drivers of tumorigenesis and have been associated with drug response and prognosis in human patients ^11-16^. Despite the importance of CNAs in cancer, they are rarely characterized in PDX models, and comprehensive analysis of CNA dynamics during *in vivo* PDX passaging has yet to be reported ^6,8,17,18^.

Here, we systematically analyze landscapes of aneuploidy and large CNAs in PDX models across multiple human cancers. We generated a comprehensive CNA catalogue of 1,110 PDX samples from 24 cancer types, using available data from SNP arrays, CGH arrays, and gene expression profiles. We used these data to characterize CNA dynamics during PDX derivation and propagation, to study the origin of passaging-acquired CNAs, and to compare PDX genomic stability across cancer types. We also compared the CNA dynamics observed in PDX models to those of newly-derived tumor cell lines and cell line-derived xenografts (CLDXs). Finally, we compared the CNA landscapes of PDXs to those of human primary tumors. We found that despite their overall similarity, the CNA landscape of PDXs diverges substantially during passaging. We discuss the potential phenotypic implications of such divergence, including its effect on therapeutic response.

## Results

### Generating a catalogue of aneuploidy and CNAs in PDXs

To enable a comprehensive analysis of aneuploidy and CNAs in PDXs, we created an integrated CNA dataset representing 1,100 PDXs. To do this, we first assembled data from DNA-based copy number measurements across multiple PDX passages, using published SNP arrays, CGH arrays and DNA sequencing data. Unfortunately, such DNA copy number data were only available for 177 PDX samples from 5 studies – too few to support a comprehensive analysis of CNA stability (**Supplementary Table 1**) ^6,7,17-19^. In contrast, gene expression profiles were available for 933 PDX samples collected from 511 PDX models across 17 studies (**Supplementary Table 1**) ^3,5,6,17,18,20-31^. To reconstruct chromosomal aneuploidy and large (>5 Mb) CNAs from these expression profiles, we used previously described computational inference algorithms that could accurately identify CNAs based on the coordinated gene expression changes induced by them ^32-34^ (**Methods**). Our final dataset thus comprised CNA data of 1,110 PDX samples from 543 unique PDX models across 24 cancer types (**Fig. 1a** and **Supplementary Data 1 and 2)**. For 342 of these PDX models, there were available data from both the primary tumor and its derived PDX model(s), or from multiple PDX passages, thus enabling an analysis of tumor evolution (**Fig.1a** and **Supplementary Table 1**).

**Figure 1:**
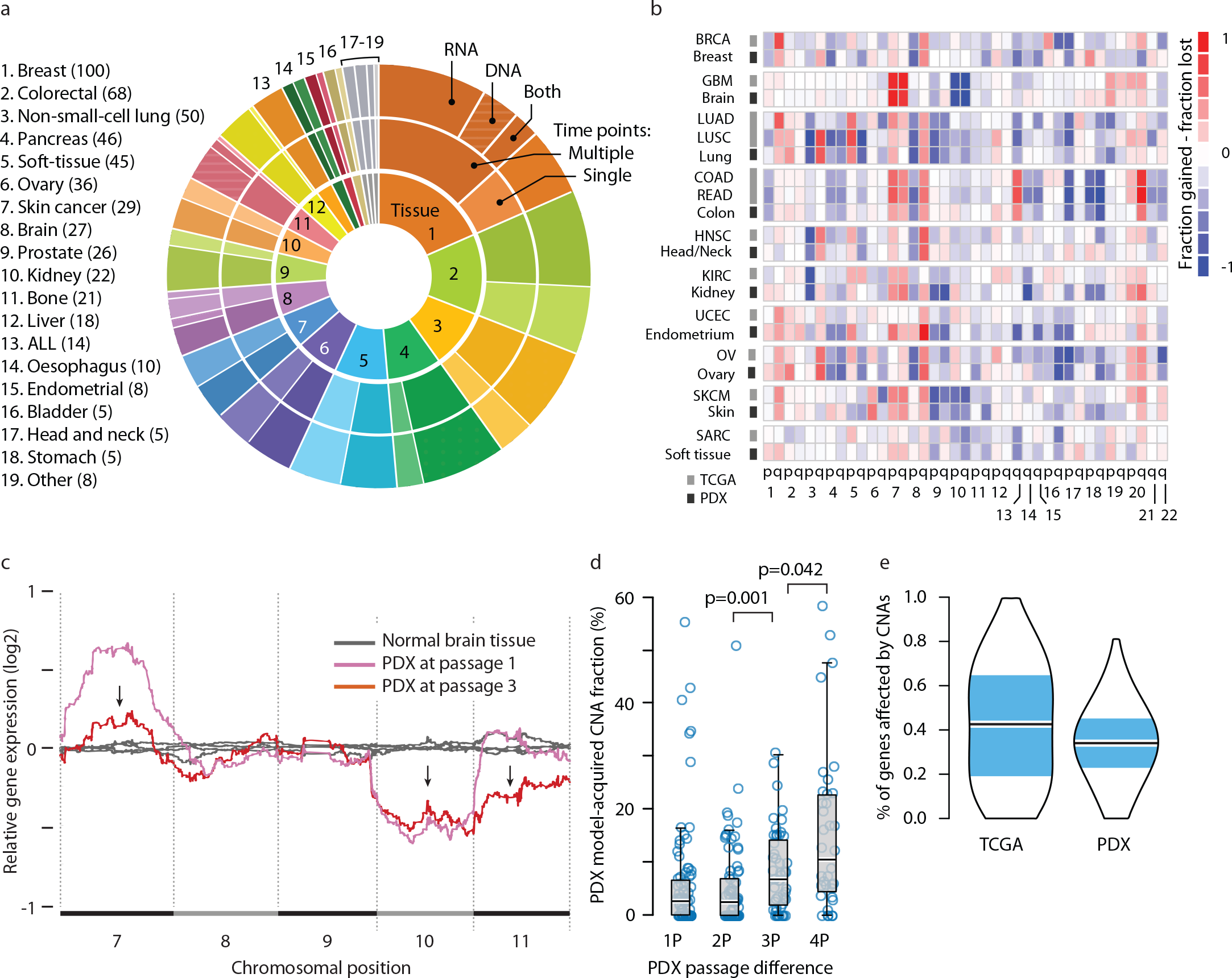
The landscape of aneuploidy and copy number alterations in PDXs. **(a)** Distribution of cancer types in our PDX dataset (n=543 unique models). In the inner circle models are divided by their lineage: each cancer type is denoted by a color and a number. In the middle circle models are divided by the number of time points analyzed: multiple time points are denoted by a darker color, and enable to follow PDX evolution throughout *in vivo* propagation. In the outer circle models are divided by the biological material from which CNAs were inferred: DNA (stripes), RNA (dots) or both (stripes and dots). (**b**) A heatmap comparing the landscapes of lineage-matched arm-level CNAs of PDXs and of primary TCGA tumors, showing an overall high degree of concordance (mean Pearson’s r = 0.79). The color of each chromosome arm represents the fraction difference between gains and losses of that arm. (**c**) A representative example of PDX model evolution. Shown are gene expression moving average plots of normal brain tissue (gray), GBM PDX model at p1 (pink) and GBM model at p3 (red), revealing the disappearance of trisomy 7, the retention of monosomy 10, and the emergence of monosomy 11, within two *in vivo* passages. (**d**) Gradual evolution of CNA landscapes throughout PDX passaging. Box plots present model-acquired CNA fraction as a function of the number of passages between measurements. Bar, median; box, 25^th^ and 75^th^ percentiles; whiskers, data within 1.5*IQR of lower or upper quartile; circles: all data points. P-values indicate significance from a Wilcoxon rank-sum test. (**e**) Violin plots present the proportion of genes affected by CNAs in TCGA and in PDX tumor samples (all tissue types combined), showing an overall similarity between both datasets. Bar, median; colored rectangle, 25^th^ and 75^th^ percentiles; width of the violin indicates frequency at that CNA fraction level.

To validate the accuracy of inferred CNAs, we analyzed PDXs from which both gene expression and SNP array data (a more direct measurement of DNA copy number) were available. Because in most cases DNA and RNA were obtained from different PDX passages, we focused on the 59 PDX models that had stable CNAs over time. The DNA- and RNA-derived profiles were highly concordant both when comparing the proportion of the genome affected by CNAs (Pearson’s r = 0.86) and when comparing the concordance of affected genes (median concordance = 0.82) (**Supplementary Figure 1**). Moreover, for 15 breast and prostate PDXs, we could directly compare the changes that occurred during their engraftment and/or passaging (hereinafter called ‘model-acquired CNAs’), using DNA and RNA data from the same samples. These DNA- and RNA-derived profiles were highly concordant (Pearson’s r = 0.95; median concordance = 0.91). These results thus confirmed that gene expression accurately identifies model-acquired CNAs (**Supplementary Table 2**).

The CNA landscapes of PDXs in our analysis were highly similar to the CNA landscapes of their respective tumor types in The Cancer Genome Atlas (TCGA) (mean Pearson’s r = 0.79; **Fig. 1b** and **Supplementary Fig. 2**), consistent with prior reports ^6,8,17,18^. This confirms that PDX models are generally genomically representative of primary tumors.

### Tracking CNA dynamics during PDX derivation and propagation

We next set out to follow CNA dynamics in individual PDX models, in order to assess their stability as well as their similarity to the tumors from which they were derived. For each model, the earliest passage (in most cases, P0 or P1) was compared to later passages in order to determine the changes that occurred throughout passaging. A representative example of PDX model evolution is shown in **Fig. 1c**.

We found that large (>5Mb) CNAs arose in PDXs rapidly: 60% of the PDX models acquired at least one large chromosomal aberration within a single *in vivo* passage, and 88% acquired at least one large aberration within four passages (**Supplementary Fig. 3a**). The CNA landscape of PDX models thus gradually shifted away from that of the original primary tumors, with a median of 12.3% of the genome (range, 0% to 58.8%) affected by model-acquired CNAs within four passages (**Fig. 1d**). Of note, similar results were obtained using three different definitions of CNA prevalence: the proportion of the genome affected by CNAs (CNA fraction), the number of discrete events, or the proportion of altered genes (**Fig. 1d** and **Supplementary Fig. 3b, c**), thereby highlighting the robustness of this finding.

There was no significant change in the overall number of CNAs throughout passaging (**Supplementary Fig. 3d**), indicating equal rates of acquiring new events and losing existing ones. We found that a median of 35.6% of the genome was affected by CNAs, consistent with prior estimates in primary tumors ^9^ (**Fig. 1e** and **Supplementary Fig. 3c**). The disappearance of CNAs during passaging was not due to changes in tumor sample purity (for instance, contamination with mouse tissue might dilute the CNA signal), as other primary events were readily detected at similar signal strength. Importantly, approximately one out of six large CNAs identified in PDX models at passage 4 was not observed in the primary tumor from which they were derived. A similar proportion of primary clonal CNAs could no longer be detected in PDXs by passage 4. We conclude that individual PDX models can quickly genomically diverge from their parental primary tumors.

### Selection of pre-existing subclones underlies CNA dynamics

Our observation that CNAs were often gained or lost during PDX passage might be explained by expansion of pre-existing subclones, the acquisition and expansion of *de novo* events, or a combination of both. Several lines of evidence suggest that clonal selection of preexisting subclones plays a major role in shaping the CNA landscape of PDXs. First, most model-acquired CNAs occurred at the earliest passages: while CNAs accumulated with each passage, the rate of their acquisition decreased over time (**Fig. 2a**). Second, gene expression signature scores indicate that apoptosis gradually decreases and proliferation increases with PDX passage number, in line with clonal selection of fitter clones (**Fig. 2b**). Third, previous analyses of clinical samples reported metastases to be more aneuploid and more chromosomally unstable on average than primary tumors ^14^. Therefore, if model-acquired CNAs were predominantly the result of genomic instability (rather than clonal dynamics), metastasis-derived PDX models should acquire more CNAs during their *in vivo* passaging. In contrast to this prediction, while we found that PDX models from metastases were indeed more aneuploid than those from primary tumors (**Supplementary Fig. 4a**) and exhibited higher chromosomal instability signature scores (**Supplementary Fig. 4b**), we did not find an elevated rate of model-acquired CNAs in metastasis-derived PDXs (**Fig. 2c**).

**Figure 2:**
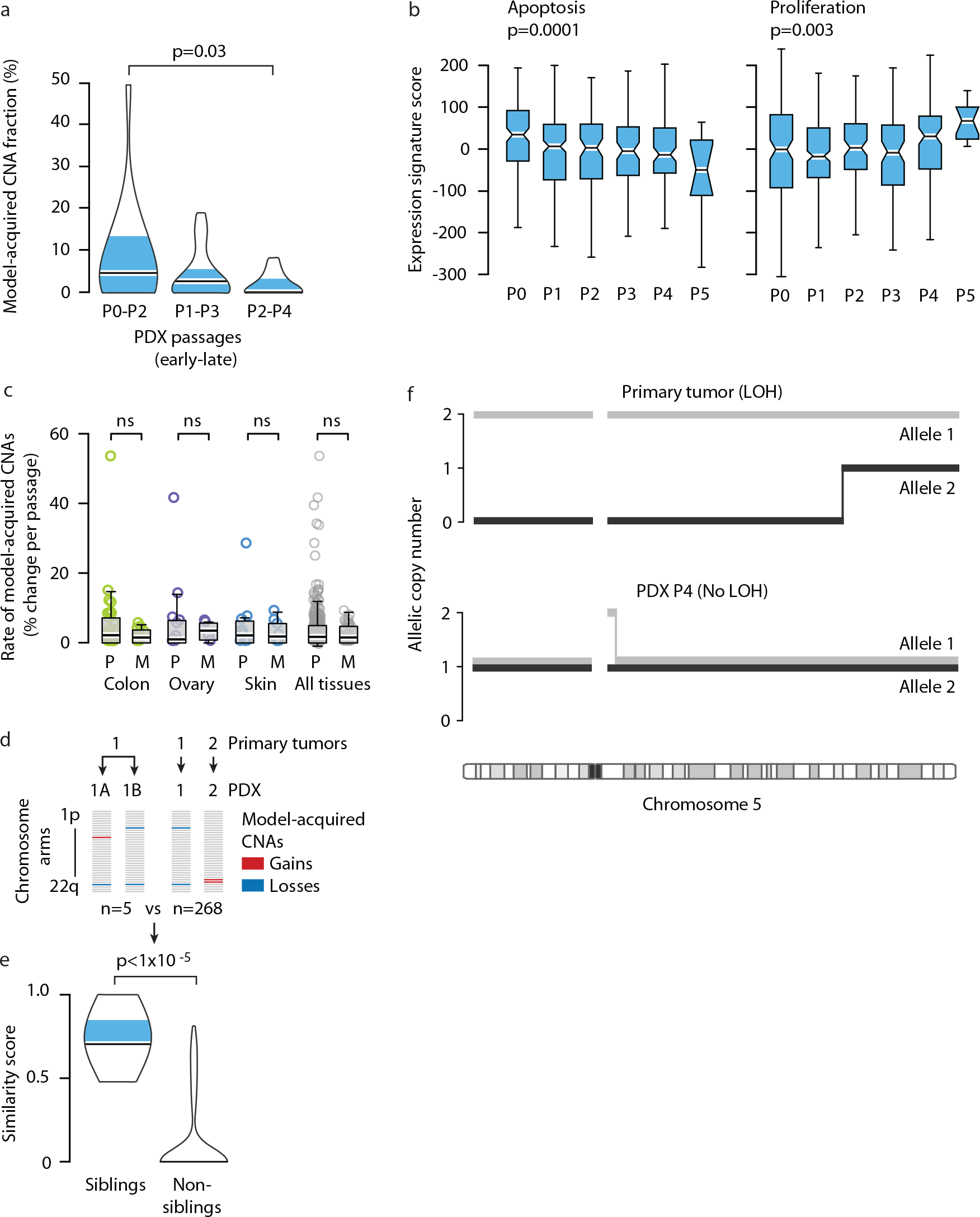
Selection of pre-existing subclones underlies CNA dynamics. **(a)**. The rate of model-acquired CNAs decreases with PDX passaging. Violin plots present the fraction of CNAs acquired within two *in vivo* passages as a function of passage number. P-value indicates significance from a Wilcoxon rank-sum test. (**b**) Apoptosis decreases and proliferation increases with PDX passaging. Box plots present the apoptosis (left panel) and proliferation (right panel) gene expression signature scores as a function of passage number. P-values indicate significance from a Kruskal-Wallis test. (**c**) Similar CNA acquisition rates in PDXs from primary tumors and from metastases. Box plots present the rate of model-acquired CNAs as a function of tumor source (P=primary, M=metastasis), across three available tissue types. n.s., non-significant (Wilcoxon rank-sum test). (**d**) Schematics showing the calculation of pair-wise similarity scores for PDX models coming from the same primary tumor but propagated independently in the mouse (“sibling” PDXs; n=5) and for PDX models coming from distinct primary tumors (“non-sibling” PDXs; n=268). (**e**) “Sibling” PDXs tend to acquire more similar aberrations than lineage-matched “non-sibling” PDXs. Violin plots present the similarity scores of “sibling” and “non-sibling” PDXs. P-value indicates significance from a lineage-controlled permutation test. (**f**) Alleles that seem to have been lost in primary tumors can “re-appear” in PDXs, demonstrating expansion of rare pre-existing subclones throughout PDX propagation. Plots present the copy number of both of chromosome 5 alleles in a primary tumor and its derived PDX. Loss of heterozygosity (LOH) is identified in the primary tumor along most of chromosome 5, but both alleles are detected in a 1:1 ratio in the PDX derived from that primary tumor.

If our hypothesis that acquired CNAs were the result of positive biological selection of existing, minor subclones, then one would expect that the same minor clones would be enriched in multiple independent grafts of the same tumor(i.e., transplanted into different “sibling” P0 mice). Five such PDX pairs (representing breast, lung, pancreas and skin cancer PDXs) were available for analysis to determine whether more model-acquired CNAs were observed in common than expected by chance (**Fig. 2d**). In principle, such clonal dynamics could arise from a bias due to selective pressure toward *de novo* events that enhance successful engraftment. To control for this, the analysis took into account the baseline frequency of each event in PDXs of the same tissue type (**Methods**). The observed similarity in model-acquired CNAs between “sibling” PDXs was significantly higher than the similarity between lineage-controlled “non-sibling” PDXs (p<1E-5; **Fig. 2e**). This finding suggests directional, deterministic selection of pre-existing subclones, which is consistent with observations in breast and hematopoietic cancers ^7,35^.

To further test the hypothesis that PDXs undergo expansion of minor subclones *in vivo*, we turned to an analysis of loss of heterozygosity (LOH) events. Because LOH is an irreversible event, an observation of heterozygosity at a region that exhibited LOH earlier on can only be explained by expansion of cells that did not undergo LOH in the first place. To examine whether LOH “reversion” ever occurs in PDX models, we queried copy number architecture in previously published whole-genome sequencing data from 15 breast cancer pairs of primary tumors and PDXs ^7^. We identified five cases of LOH “reversion” (**Fig. 2f** and **Supplementary Fig. 5**). For example, while the primary tumor SA494 had a clonal copy-neutral LOH throughout most of chromosome 5 (allelic ratio ~1 for the major allele and ~0 for the minor allele), the PDX model derived from that tumor reverted to heterozygosity by passage 4 (allelic ratio ~0.5 for both alleles) (**Fig. 2f**). These analyses thus confirm that rare pre-existing subclones that are not readily detected in a population-level analysis of the primary tumor, can expand and become the dominant clone in PDXs within as little as one *in vivo* passage.

We therefore conclude that CNA dynamics are strongest during engraftment and the first few *in vivo* passages, continue at a reduced rate throughout model propagation, and most likely result to a large extent from selection of pre-existing subclones.

### The degree of genomic instability in PDXs mirrors that of primary tumors

As human cancer types differ considerably in their CNA prevalence and rate of acquisition (also referred to as degree of genomic instability, or DGI), we next compared CNA dynamics in PDXs across cancer types. We found that the rate of model-acquired CNAs varies significantly (p=0.001 comparing the most stable to the most unstable tumor types), with brain tumor PDXs being the most stable and gastric tumors being the most unstable (a median of 0% and 4.2% of the genome affected by CNAs per passage, respectively) (**Fig. 3a**).

**Figure 3:**
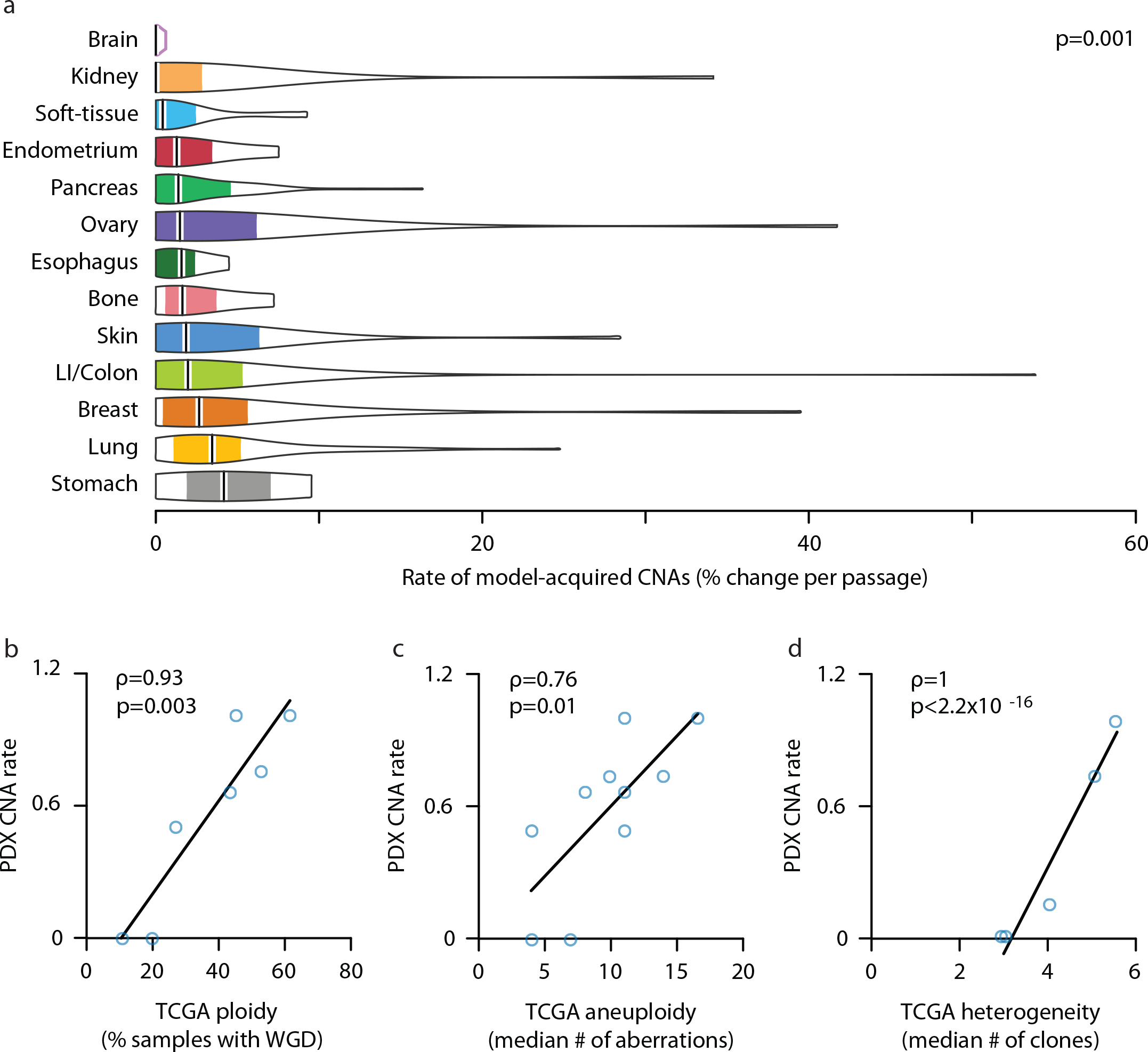
Genomic instability of PDXs mirrors that of primary tumors. **(a)** The degree of genomic instability (DGI) of PDXs is cancer type-specific. Violin plots present the rate of CNA acquisition throughout PDX propagation of 13 cancer types. P-value indicates significance from a Wilcoxon rank-sum test. (**b**) The DGI of PDXs and that of primary tumors correlate extremely well. In PDXs, tissue DGI was defined as the median number of CNAs per passage. In TCGA tumors, tissue DGI was defined as the fraction of samples with whole-genome duplication (WGD). (**c**) This correlation holds when the tissue DGI is defined, both for PDXs and for TCGA tumors, by the median number of arm-level CNAs. (**d**) The DGI of PDXs also correlates extremely well with intra-tumor heterogeneity (ITH) of primary tumors (excluding the skin tissue). The DGI of PDXs was defined as the median number of arm-level CNAs per passage. The heterogeneity of primary tumors was defined as the median number of clones per tumor. Spearman’s rho values and p-values indicate the strength and significance of the correlations, respectively.

We therefore asked whether this spectrum of PDX aneuploidy was reflective of the aneuploidy levels of human cancer types. We measured aneuploidy in TCGA data according to two metrics. First, we used the previously reported percentage of samples with whole-genome duplication (WGD) ^9^. Across the seven tissues for which data were available from both TCGA and PDX datasets, the CNA acquisition rate in PDX samples correlated strongly with WGD prevalence in TCGA samples, whether PDX rate was evaluated by CNA fraction (Spearman’s rho = 0.88, p=0.085; **Supplementary Fig. 6a**) or by the number of discrete events (Spearman’s rho = 0.93, p=0.003; **Fig. 3b**). Second, we found that the median number of model-acquired arm-level CNAs in PDXs and the median number of arm-level events acquired during tumor development in TCGA samples correlated well across 10 different cancer types (Spearman’s rho = 0.76, p=0.010; **Fig. 3c**). We thus conclude that the DGI variation among PDX tumors types represents that of the primary tumors.

This association between DGI in PDX models and human cancers of the same tissue type may result from similar rates of ongoing acquisition of new events, or from increased levels of intra-tumor heterogeneity in highly aneuploid tumors ^16^. As we found that clonal selection/drift of pre-existing events had a major role in shaping the CNA landscape of PDXs, we examined whether the tissue-specific rate of CNA dynamics correlates with the degree of heterogeneity that characterizes each cancer type. The CNA acquisition rate in PDXs correlated well with the median number of clones of the respective primary tumor type ^16^, across the six cancer types that could be matched (Spearman’s rho = 0.82, p=0.044; **Supplementary Fig. 6b**). Interestingly, melanoma had the highest degree of intra-tumor heterogeneity, but only a moderate level of DGI in PDXs, and was therefore the only cancer type that significantly deviated from the observed correlation; the correlation became even stronger when melanoma was removed from the analysis (Spearman’s rho = 1, p<2.2E-16; **Fig. 3d**). A potential explanation for this discrepancy is that melanoma is the only cancer type for which subcutaneous injection is very similar to orthotopic injection, potentially leading to weaker differences in selection pressures between the human and mouse environments. Alternatively, the extent of genetic heterogeneity in melanoma may be overestimated compared to other cancer types due to the unusually high mutation load of this tumor.

The combined results of these analyses suggest that PDX models have characteristic tissue-specific levels of CNA dynamics, which correspond both to the DGI and to the degree of heterogeneity of the respective primary tumor types. As genetic heterogeneity is closely associated with aneuploidy levels and DGI in primary tumors ^16,36,37^, either of these factors – or both of them together – could explain the observed correlations.

### CNA recurrence analysis reveals distinct selection pressures in PDXs vs. primary tumors

A key question is whether the clonal dynamics observed in PDXs mimic the selective pressures observed in human patients. If so, then the clonal dynamics of PDXs could serve as a useful model for the evolution of human tumors; if not, then PDX-acquired genetic events would gradually shift them away from the human tumors from which they were derived, as a result of murine-specific selective pressure. To address this, we asked whether recurrent arm-level genetic events that are observed in human tumors remain under selective pressure when transplanted into mice; loss of these signature events would signal significant differences in selective pressures between human and mouse hosts.

To test this, we identified 61 recurrent arm-level CNAs across TCGA tumor types, and followed these recurrent CNAs in PDXs. Surprisingly, events that were recurrent in the TCGA dataset tended to disappear throughout PDX passaging. Specifically, among lineage-matched PDXs, we observed 116 model-acquired events that were in the opposite direction to the recurrent TCGA CNAs, and only 79 model-acquired events in the same direction (p = 0.01, McNemar’s test). We identified twelve recurrent events in TCGA samples across five cancer types (GBM, breast, lung, colon and pancreatic cancer) which were preferentially lost throughout PDX passaging (**Fig. 4a** and **Supplementary Fig. 7**). Events that tend to disappear throughout PDX propagation should be less prevalent at high passage compared to low passage PDXs of the respective lineage. Indeed, nine of the twelve events that PDXs tend to lose, including the hallmark gains of chr1q and 8q in breast cancer and chr7 in GBM, and the hallmark losses of chr10 in GBM and chr4q in non-small cell lung cancers, were less common in high passage PDXs (**Fig. 4b** and **Supplementary Fig. 8**), although in only three cases did this reach statistical significance, likely reflecting the small sample size of each group.

**Figure 4:**
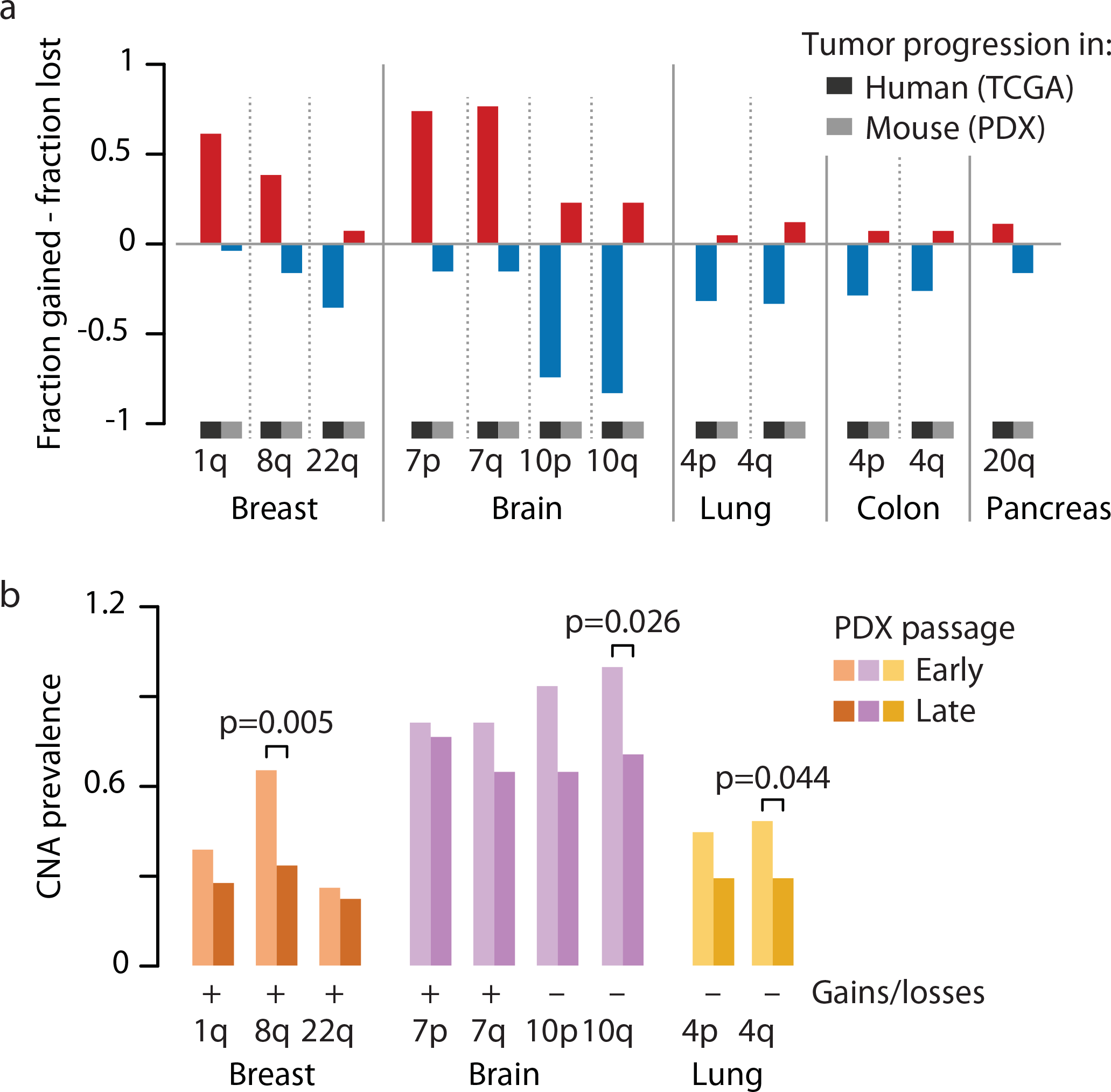
Tumor evolution of PDXs diverges from that of primary tumors. **(a)** Opposite propensities to gains and losses in human tumors and PDX models. Bar plots present the fraction difference between gains and losses of 12 recurrent TCGA arm-level CNAs. The PDX fractions represent the model-acquired CNAs, rather than the absolute prevalence of these events. (**b**) Recurrent TCGA arm-level CNAs are more common in early passage PDXs than in late passage PDXs. Bar plots present the absolute prevalence of each event in the relevant cancer type. P-values indicate significance from a Fisher’s exact test.

Together, these data demonstrate that PDXs can lose recurrent chromosomal aberrations that may play causal roles in the development of tumors in humans. This suggests that the selection pressures that led to the acquisition and retention of these hallmark CNAs in patients may no longer apply to the murine model environment.

### Genomic instability of PDXs is comparable to that of cell lines and CLDXs

PDXs are generally considered to more faithfully reflect primary human tumors compared to cell lines because cell lines grow in the absence of signals from the tumor microenvironment^1,38^. However, the immunodeficient, subcutaneous murine microenvironment also differs considerably from the natural human host. To address the assumption that PDXs better preserve the fidelity of human tumors, we analyzed the CNA dynamics of PDXs *in vivo* compared to those of cell lines *in vitro*.

We found that the prevalence of model-acquired CNAs is similar in newly-derived cell lines compared to PDXs. We analyzed the CNA landscapes of nine new cell lines derived in our lab from five cancer types (colon, GBM, pancreas, esophagus and thyroid) (Tseng et al., manuscript in preparation; **Supplementary Table 3**). These cell lines were subjected to whole exome sequencing at four or five time points throughout their propagation, from p0/p1 to p20, and the CNA landscape of each sample (n=38) was determined (**Methods**). Similar to our observations with PDX models, newly-derived cell lines acquired CNAs with passaging, and their CNA landscape gradually shifted away from that of the earliest passage (**Supplementary Fig. 9a**). As seen in PDXs, most of the changes occurred during the first few passages, and the rate of model-acquired CNAs decreased throughout culture propagation (**Fig. 5a**). Notably, while CNA rates (defined as the fraction of the genome affected by model-acquired CNAs per passage) varied considerably among cell lines, they fell well within the range seen in PDXs, in a lineage-matched comparison (p=0.55; **Fig. 5b**). These results from our new cell line models were recapitulated with newly-derived cell lines in three independent studies of GBM, kidney and head and neck cancer (n=31; **Supplementary Fig. 9b** and **Supplementary Table 3**), suggesting that they do not depend on any particular cell line propagation method.

**Figure 5:**
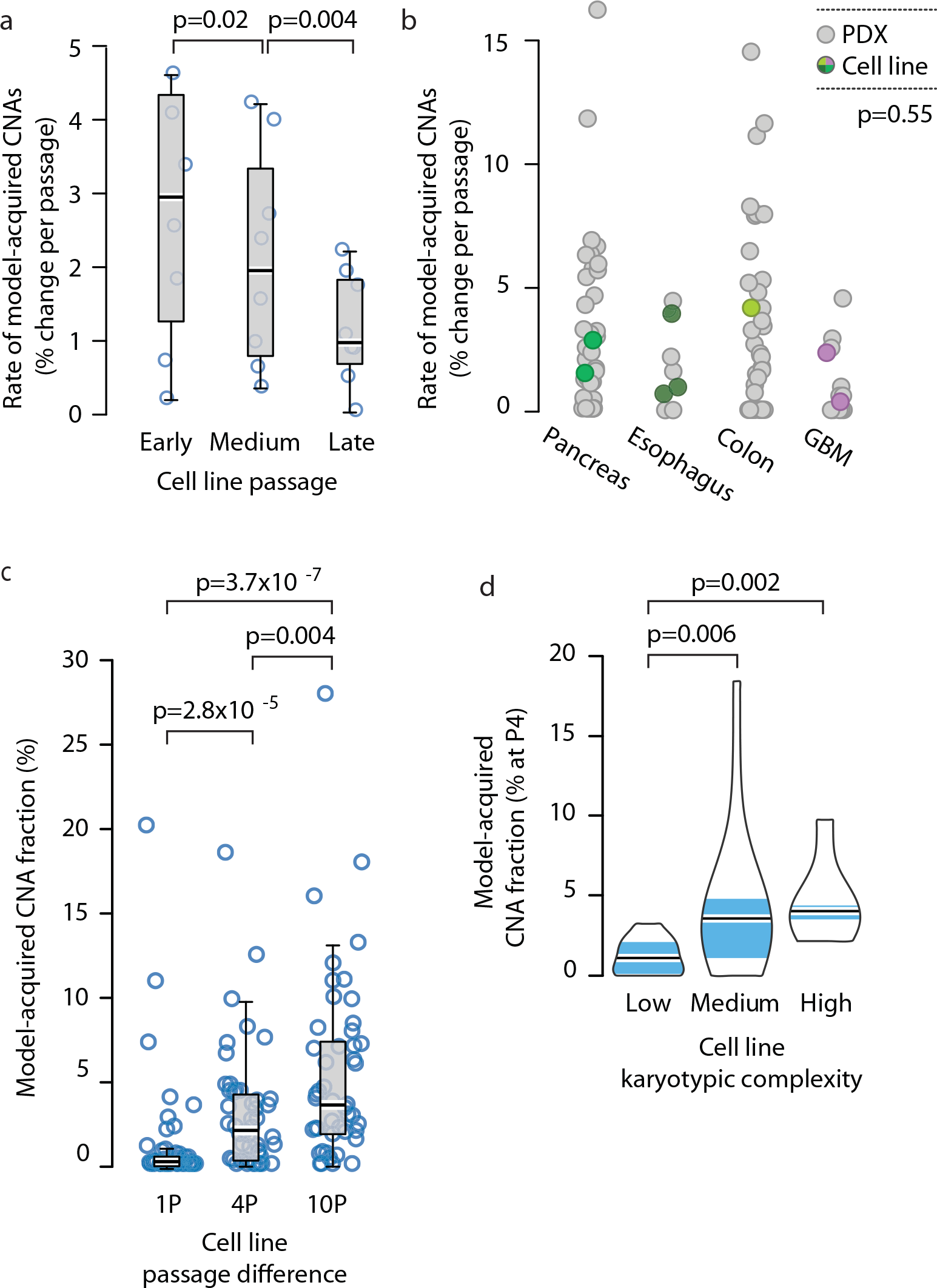
Genomic instability of PDXs is comparable to that of cell lines and CLDXs. (**a**) The rate of CNA acquisition decreases with cell line passaging. Box plots present the rate of CNA acquisition as a function of *in vitro* passage number. P-values indicate significance from a Wilcoxon rank-sum test. (**b**) Similar rates of CNA acquisition in PDXs and in newly-derived cell lines. Dot plots present the distribution of model-acquired CNA rates across four available cancer types. P-value indicates lack of significance from a lineage-controlled permutation test. (**c**) Gradual evolution of CNA landscapes throughout CLDX passaging. Box plots present model-acquired CNA fraction as a function of the number of passages between measurements. P-values indicate significance from a Wilcoxon rank-sum test. (**d**) The CNA acquisition rate of CLDXs is associated with the numerical karyotypic complexity of the parental cell lines. Violin plots present the fraction of CNAs acquired by passage 4 as a function of numerical karyotypic complexity. P-values indicate significance from a Wilcoxon rank-sum test.

Next, we compared CNA dynamics between PDXs and cell line-derived xenografts (CLDXs). To assess CNA dynamics during the *in vivo* propagation of established cancer cell lines, we turned to the NCI MicroXeno project, which profiled gene expression of 49 well-studied, established human cancer cell lines across multiple *in vivo* passages 39. We used the same computational algorithms ^32-34^ that we applied to the PDX models to infer aneuploidy and CNAs from these gene expression profiles, resulting in 823 copy number profiles (**Supplementary Data 3 and 4**). We found that CNAs accumulate with *in vivo* passaging of CLDXs (**Fig. 5c**), and that the DGI of CLDXs correlates with the karyotypic complexity of their parental cell lines (**Fig. 5d** and **Supplementary Fig. 9d**), similar to what we observed in PDXs.

However, the rate of CNA acquisition was lower in CLDXs: within four passages, the median model-acquired CNA fraction was 2.2% in CLDXs, compared to 12.3% in PDXs (p=1.6E-6), likely reflecting the reduced heterogeneity of established cell lines compared to primary tumors at the time of xenograft initiation ^40^. Taken together, our data from three types of cancer models (PDXs, cell lines, and CLDXs) demonstrate that switching the environment in which a model is propagated results in CNA dynamics that gradually alter its CNA landscape. All cancer models are subject to such clonal selection; unfortunately, PDXs are not spared.

### CNA dynamics in PDXs can affect their drug response

It is conceivable that while PDXs undergo selection in the mouse, such selection is unimportant with respect to modeling therapeutic response. To address this, we turned to a dataset of PDXs with accompanying responses to both genotoxic chemotherapies and targeted therapeutics ^3^.

Both very low and very high levels of aneuploidy have been associated with response to genotoxic drugs and improved patient survival ^14,16,41,42^. Importantly, CNA acquisition rate (DGI), rather than absolute levels of aneuploidy, determines sensitivity to further perturbation of chromosome segregation ^15^. We therefore determined the DGI of PDX models, and asked whether it similarly predicts response to chemotherapies (**Methods**). For three of five chemotherapies tested, extreme (either very low or very high) levels of DGI – but not overall aneuploidy levels – were associated with favorable therapeutic response (**Fig. 6a**): dacarbazine in skin PDXs, paclitaxel in lung PDXs, and abraxane/gemcitabine in pancreas PDXs (p=0.04, 0.014 and 0.006, respectively). The biological activity and clinical efficacy of these drugs were previously linked to chromosomal instability ^43-47^. These results indicate that PDXs recapitulate the observations in patients that overall levels of genomic instability are correlated with response to cytotoxic chemotherapies.

**Figure 6:**
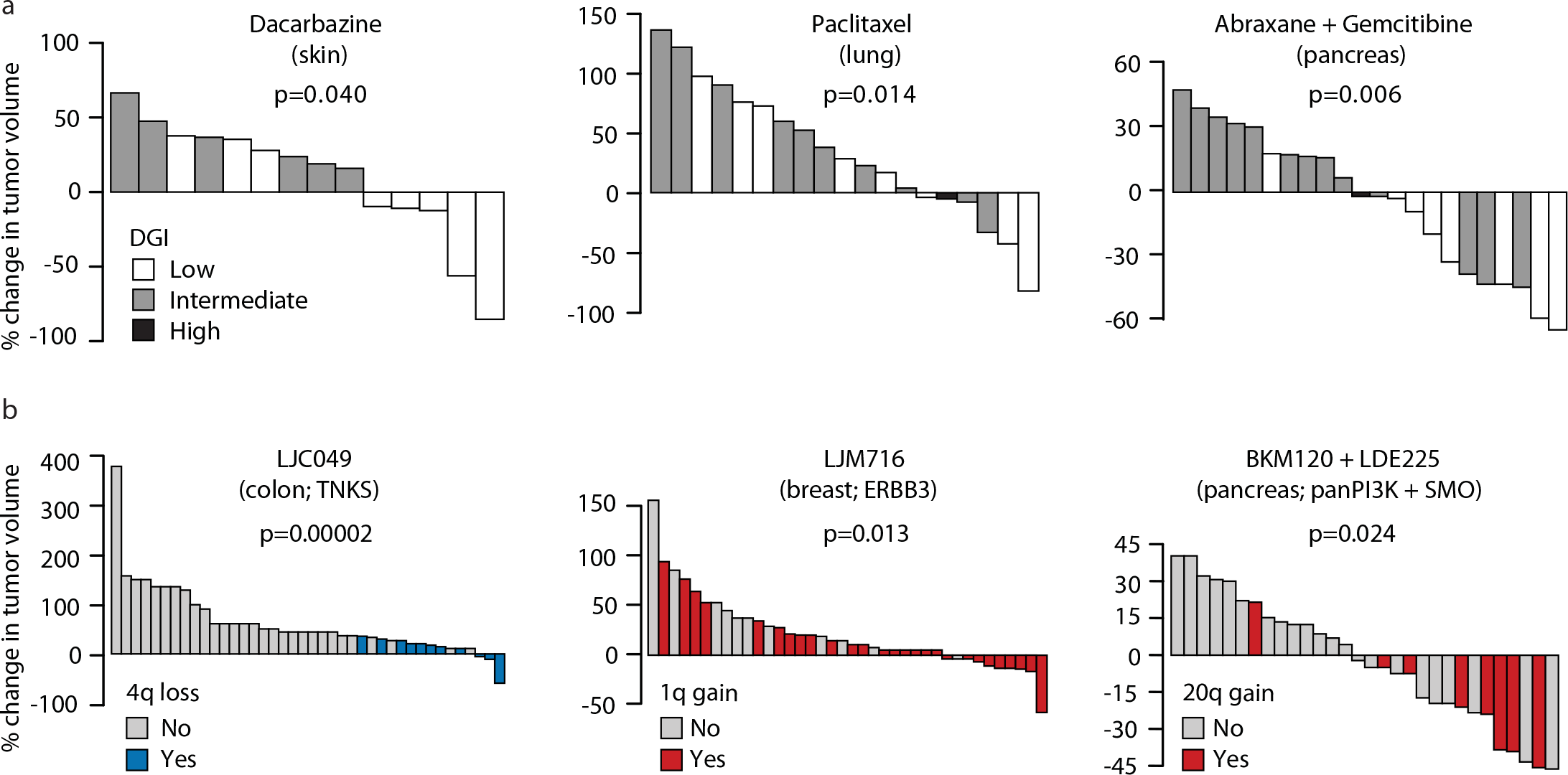
CNA dynamics affect PDX drug response. (**a**) Extreme levels of genomic instability are associated with better therapeutic response to chemotherapies. Waterfall plots present the response to dacarbazine (n=14), paclitaxel (n=19), and the combination of abraxane and gemcitabine (n=22) in skin, lung and pancreas PDXs, respectively. DGI, degree of genomic instability. P-values indicate significance from a Wilcoxon rank-sum test. (**b**) The status of recurrent arm-level CNAs is associated with response to targeted therapies. Waterfall plots present the response to the TNKS inhibitor LCJ049 (n=40), the ERBB3 inhibitor LJM716 (n=38), and the combination of the PI3K inhibitor BKM120 and the SMO inhibitor LDE225 (n=31). P-values indicate significance from a Wilcoxon rank-sum test.

We next asked whether particular model-acquired CNAs might affect PDX responses to targeted therapies, given that specific recurrent arm-level or whole-chromosome events have been reported to alter the cellular response to certain drugs ^48-50^. We evaluated the association between PDX response to targeted therapies and the presence or absence of individual arm-level CNAs, focusing on the twelve driver CNAs that we found to be selected against during PDX passaging. After correction for multiple hypothesis testing, we identified three statistically significant drug response-CNA associations (**Fig. 6b**): chromosome 4 monosomy was associated with increased response of colon PDXs to the TNKS inhibitor LCJ049 (p=0.005, q=0.04 for 4p loss, and p=0.00002, q=0.0003, for 4q loss); chromosome 20q gain was associated with increased response of pancreatic PDXs to the combination of the PI3K inhibitor BKM120 and the SMO inhibitor LDE225 (p=0.024, q=0.19); and chromosome 1q gain was associated with increased response of breast PDXs to the ERBB3 inhibitor LJM716 (p=0.013, q=0.23). These results indicate that it is not unusual for CNAs (and presumably other genomic events) that undergo negative selection in the murine host to be associated with changes in sensitivity to specific targeted agents. Such associations may have important implications for the development of predictive biomarkers based on PDX results.

## Discussion

The ability to directly transfer human tumors into mice, and propagate them for multiple passages *in vivo*, offers unique opportunities for cancer research and drug discovery, making PDXs a valuable cancer model. Like any other model system, however, understanding the limitations of the model – and the ways in which it differs from human tumors in their natural environment – is required for its optimal application. Our findings suggest that the genomic instability of PDXs has been underappreciated: the CNA landscapes of PDXs change continuously, and so their propagation distances them from the primary tumors from which they were derived.

We note that our analysis focused on CNAs because the availability of large numbers of gene expression profiles of PDX tumors allowed for the reconstruction of their CNA landscapes, thus generating a sufficiently well-powered dataset. The phenomenon of model-acquired genetic events, however, is unlikely to be restricted to CNAs. Rather, it seems likely that PDX models (as all other models) also acquire other types of aberrations, including point mutations, small insertions and deletions, translocations, and epigenetic modifications. Large-scale datasets do not exist at present to experimentally confirm these other forms of selection.

As our analysis was based on bulk-population measurements, the cellular origin of each model-acquired event could not be definitively determined. Our study strongly suggests that clonal dynamics play a major role in model-acquired CNAs, especially at the early stages of PDX derivation and propagation. In particular, the acquisition of identical events in “sibling” PDXs, and the detection of LOH “reversion” throughout PDX passaging, strongly point towards expansion of pre-existing subclones. However, our analysis suggests that *de novo* events also occur. First, model-acquired CNAs are not limited to the early passages and keep emerging, albeit at a lower rate, even at high passages (**Supplementary Fig. 10a** and **Supplementary Data 2**). Second, although “sibling” PDXs exhibit high similarity of model-acquired CNAs, most of them also acquire unique events (**Supplementary Figure 10b**). Third, we found that PDXs with a mutant or deleted p53 present a significantly higher rate of CNA acquisition throughout passaging, compared to their WT counterparts (**Supplementary Figure 10c**). All three of these findings could also be potentially explained by extensive pre-existing heterogeneity, however. Regardless of their exact origin, we found that CNAs often became fixed in the population quickly, as a single *in vivo* passage sometimes rendered a chromosomal aberration that had been completely undetected readily identified at the population level.

Unique, context-dependent selection pressures shape tumor evolution, giving rise to recurrent cancer type-specific CNAs ^9^. The strong clonal dynamics observed in PDX models at early passages, together with the tendency of recurrent arm-level events to disappear throughout PDX propagation, suggest that tumor evolution trajectories differ between patients and PDX models, most likely due to distinct selection pressures in these different environments ^2^. At least three important parameters may account for these differences: the species (human vs. mouse), the anatomical and physiological context (a specific organ vs. subcutaneous growth), and the interaction with the immune system (immunocompetent patients vs. immunodeficient animals). In the future, comparisons of orthotopic vs. subcutaneous PDXs, and of mouse-derived xenografts in “humanized” immunocompetent vs. immunodeficient recipients, may help delineate the contribution of each of these parameters to shaping tumor evolutionary pressures.

Recent genomic analyses revealed that metastases evolve independently from the primary tumors, often representing common ancestral subclones that are not detected in individual biopsies of the primary tumors. In contrast to the considerable heterogeneity between primary tumors and metastases, distinct metastatic sites tend to be relatively homogeneous ^51-53^. Our findings from PDXs echo those from studies of metastasis: the dominant clones in the PDXs often come from minor subclones of the primary tumors, and PDXs that originate from the same primary tumor (the equivalence of multiple metastatic sites) tend to evolve in similar trajectories. It has been suggested that caution is required when inferring the genetic composition of metastatic disease from a biopsy of the primary tumor, and vice versa ^51-53^; similarly, we propose that the genetic composition of a PDX tumor may differ from its primary tumor of origin, potentially in therapeutically meaningful ways.

PDX collections are generally used for drug testing in two different ways: to predict, at the cohort level, the relationship between genotype and dependency; and to predict, at the individual level, the therapeutic response ^4^. Our findings have several practical implications for both of these uses. The rapid genomic divergence that we identify on the individual tumor level suggests that PDXs may often not be faithful representations of their parental tumors. If individual PDXs are to be utilized as avatar models for personalized medicine, it will be necessary to ensure that the model retains the relevant genomic features of the primary tumor from which it was derived, before PDX drug response is used to guide clinical treatment decisions. It will also be advisable to use such avatar models at the earliest passage possible and avoid their prolonged propagation, especially in the context of a 1x1x1 (one animal per model per treatment) experimental design ^6^. For population level analyses, our findings highlight the need to document the molecular properties of the models at the same passage as that used for drug testing, rather than relying on an early passage characterization. They also emphasize the importance of large cohorts of PDX models, similar to the large cell line collections that were recently established ^54,55^, in order to average out random effects when performing drug screens and biomarker studies. Finally, the gradual loss of recurrent primary CNAs suggests that prolonged propagation could lead to under-representation of some hallmark cancer events in late passage PDX cohorts.

The comparison of PDXs to newly-derived cell lines revealed that PDXs do not necessarily capture the genomic landscape of primary tumors better than cell lines, in contrast to common belief ^1^. The definition of a passage in cell lines differs from that in PDXs, and fewer cell divisions occur between passages *in vitro*. Therefore, the similar rate of model-acquired CNAs per passage may actually reflect greater instability in cell lines. From a practical standpoint, however, the absolute passage number comparison seems appropriate. Importantly, the similar rate of changes suggest that multiple cell line models from a single primary tumor may capture more of the original CNA landscape – and more of its heterogeneity – than a single PDX model. As costs and complexities of PDX generation are generally greater than those of cell line derivation, this should be considered and balanced against the advantages of *in vivo* studies, for the specific desired applications.

The comparison of PDXs to CLDXs showed a lower CNA acquisition rate in CLDXs than in PDXs. There are three potential explanations for this difference: a lower degree of heterogeneity in established cancer cell lines, a reduced bottleneck upon cell line transplantation, or a reduced rate of ongoing instability. As cell lines are generally more clonal than primary tumors ^40^, and as we could attribute much of the CNA dynamics observed in PDXs to expansion of pre-existing clones, we speculate that the reduced heterogeneity of established cell lines explains most of the observed difference, although this question remains to be addressed experimentally. Regardless of its source, however, this difference suggests that although the genomic landscapes of established cell lines don’t represent primary tumors as faithfully as newly-derived cell lines and PDXs, these landscapes are more stable.

Our study may have implications beyond cancer model systems. Recent single cell RNAseq studies used hallmark arm-level CNAs as genetic markers to distinguish between tumor and non-tumor cells ^56,57^. The finding that some of these events, such as trisomy 7 and monosomy 10 in GBM, can disappear in PDXs, suggests that minor subclones without these aberrations probably exist in primary tumors; therefore, cells should not be classified as non-tumor cells solely based on the absence of a single hallmark event.

In summary, we observed extensive clonal dynamics in PDXs through the analysis of aneuploidy and CNAs, and demonstrated their potential sources and implications. The same clonal dynamics should also affect additional genomic features, such as point mutations ^7,8^, for which high degrees of intra-tumor heterogeneity have also been observed in primary tumors ^16,38^. Our study raises the possibility that the distinct selection pressures in patients and in PDX models may shape the tumor mutational landscapes in different directions; such context-specific selective pressures should be considered when using experimental models to advance precision cancer medicine.

## Online Methods

### PDX data assembly and processing

CGH array, SNP array and gene expression microarray data were obtained from the GEO (http://www.ncbi.nlm.nih.gov/geo) and EMBL-EBI (http://www.ebi.ac.uk) repositories. RNA sequencing data were obtained from the NCBI Sequence Read Archive (https://www.ncbi.nlm.nih.gov/sra/). Accession numbers are provided in Supplementary Table 1. Normalized matrix files were downloaded, and samples were curated manually to identify the cancer tissue type and the PDX passage number. Arrays were analyzed for quality control and outliers were removed. The final database consisted of 1,100 PDX tumor samples, from 543 unique PDX models across 24 cancer types. The analysis was performed in batches, and normal tissue samples included in each study served as internal controls, whenever available. Data were processed using the R statistical software (http://www.r-project.org/). For all platform types, probe sets were organized by their chromosomal location, and log2-transformed values were used. Probe sets without annotated chromosomal location were removed. For gene expression data, all the probe sets of each gene were averaged (as well as their chromosomal location), in order to obtain one intensity value per gene. A threshold expression value was set, and genes with lower expression values were collectively raised to that level: flooring values were 6-7 for the Affymetrix and Illumina platforms, and −0.5 for the Agilent platforms. Probe sets not expressed in >20% of the samples within a batch were removed. The 10% of the probe sets with the most variable expression levels were also excluded, to reduce expression noise.

### Generation of CNA profiles

CNA profiles from SNP arrays were generated using the Copy Number Workflow of the Partek Genomics Suite software (http://www.partek.com/pgs), as reported by the original studies. CNA profiles from CGH arrays were generated using the CGH-Explorer software (http://heim.ifi.uio.no/bioinf/Projects/CGHExplorer/), using the program’s piecewise constant fit (PCF) algorithm, with the following set of parameters: Least allowed deviation = 0.3; Least allowed aberration size = 30; Winsorize at quantile = 0.001; Penalty = 12; Threshold = 0.01. CNA profiles from gene expression data were generated using the protocols developed by Ben-David et al ^32^ and by Fehrman et al ^34^. For all gene expression platforms, the e-karyotyping method was applied ^32^: whenever normal tissue samples were available, the median expression value of each gene across the normal samples was subtracted from the expression value of that gene in the tumor samples, in order to obtain comparative values. These relative gene expression data were then subjected to a CGH-PCF analysis, with the following set of parameters: Least allowed deviation = 0.25; Least allowed aberration size = 30; Winsorize at quantile = 0.001; Penalty = 12; Threshold = 0.01. For Affymetrix gene expression platforms, Human Genome U133A and U133Plus2.0, the functional genomic mRNA profiling (FGMP) method ^34^ was also applied: gene expression data were corrected for the first 25 previously-identified transcriptional components, and the corrected data were subjected to the same processing steps and CGH-PCF analysis described above, with the following set of parameters: Least allowed deviation = 0.15; Least allowed aberration size = 30; Winsorize at quantile = 0.001; Penalty = 12; Threshold = 0.01. CNA profiles from DNA sequencing were obtained in a processed table form from the publication by Eirew et al. ^7^. CNA profiles from SNP arrays were obtained in a processed form from the publication by Gao et al. ^3^ and compared to CNA profiles from RNA sequencing of the same PDX models. For visualization purposes, moving average plots were generated using the CGH-Explorer moving average fit tool.

### Identification of model-acquired CNAs

To identify CNAs emerging during the generation and propagation of PDXs, 342 PDX models in which data were available from multiple time points were analyzed. These PDX models were compared either to the primary tumors from which they were derived, or to their earliest available passage. For gene expression data, model-acquired CNAs were identified by ekaryotyping. For each probe set, a relative value was obtained by subtracting the early time point value from the late time point value. CGH-PCF analysis was then performed, with the same parameters described above. For visualization purposes, moving average plots were generated using the CGH-Explorer moving average fit tool.

### CNA recurrence analysis

For each tissue type, the arm-level CNA recurrence was computed and compared between the PDX dataset and the human patient TCGA dataset (http://cancergenome.nih.gov). Chromosome arm-level events in TCGA samples were called using a novel approach to be described in Taylor, Shih, Ha et al. (manuscript in preparation). Briefly, segments of CNAs identified by ABSOLUTE ^58^ were determined as loss, neutral, and gain relative to each sample’s predicted tumor ploidy. Consecutive segments were iteratively joined such that the combined segment is no less than 80% altered in a given direction (i.e. gain or loss, not both). For every combination of arm/chromosome and direction of alteration within each TCGA tumor type, the start coordinates, end coordinates, and proportion of chromosome arm altered (based on the longest joined segment) were clustered across samples using a 3-dimensional Gaussian Mixture Model. The optimal clustering solution was chosen based on the Baysian information criterion. Clusters whose mean fraction altered in either specific direction was >=80% were considered “aneuploid”, those whose mean fraction altered (in both directions) was <=20% were considered “non-aneuploid”. Chromosome arm-level events in PDX samples were determined using the CNA status of the largest overlapping segment from the e-karyotyping analysis. Chromosome arm-level events in PDX samples were called using the CNA status of the largest overlapping segment from the e-karyotyping analysis. The comparisons of absolute CNA landscapes were performed using the FGMP-derived CNA profiles, and the comparisons of model-acquired CNAs were performed using the e-karyotyping-derived CNA profiles. The comparisons between early and late passage PDXs were performed using FGMP-derived CNA profiles: samples from p<=1 were defined as early passage, and samples from p>=3 were defined as late passage. Heatmaps were generated using the ‘pheatmap’ R package, and clustering was performed using euclidean distance and complete linkage.

### DGI comparison across passages and tissue types

The degree of genomic instability (DGI) of each sample was determined in three ways: 1) the fraction of the genome affected by model-acquired CNAs per passage, 2) the number of discrete events per passage, and 3) the fraction of altered genes per passage. For each cancer type, the median number of model-acquired arm-level CNAs across all PDX samples was determined, and compared to several TCGA statistics: the percentage of samples with whole-genome duplication taken from the publication by Zack et al. ^9^, the median number of arm-level CNAs per sample, and the median number of clones per sample taken from the publication by Andor et al. ^16^.

### Similarity analysis

PDX samples derived from the same primary tumors, but propagated in different animals starting from their initial transplantation (i.e., transplanted into different P0 mice) were defined as “siblings”. PDX samples derived from the same primary tumors, and propagated in the same animal at some point during PDX propagation were excluded from the analysis. PDX samples from distinct primary tumors were defined as “non-siblings”. Similarity scores were calculated for each pair of samples, based on the arm-level events that occurred during their *in vivo* passaging (i.e., model-acquired CNAs), using a modified Jaccard similarity coefficient. This similarity coefficient was inversely weighted to account for the observed prevalence of each CNA in each PDX tissue type. Therefore, the similarity score was calculated using the following equation: 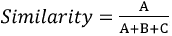, where 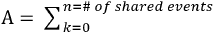1 freq(k), 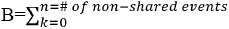 1, 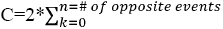 1, and freq (k) is the frequency of event k in that tumor type.

### Loss of heterozygosity analysis

Allelic copy number data were obtained from the publication by Eirew et al. ^7^. Using 10Mb windows along the genome, we identified the following scenarios: (1) the minor allele was 0 (LOH) in a primary tumor but > 0 (presence of both alleles) in the tumor-derived PDX model, and (2) the minor allele was 0 (LOH) in an early passage PDX but > 0 in a later passage of the same PDX model. These instances of apparent “reversion” of LOH were visualized using the Integrative Genomics Viewer (IGV; https://www.broadinstitute.org/igv/) and re-plotted in Figure 2f.

### Gene expression signature scores

The apoptosis and proliferation gene sets were derived from the Molecular Signature Databse (http://software.broadinstitute.org/gsea/msigdb), using the “Hallmark_Apoptosis” ^59^and the “Benporath_Proliferation” ^60^ gene sets, respectively. The CIN70 gene set was derived from the publication by Carter et al. ^13^. Signature scores were generated for all PDXE models ^3^. For each gene set, genes not expressed at all in the PDX dataset were removed, and the remaining gene expression values were log2-transformed and scaled by subtracting the gene expression means. The signature score was defined as the sum of these scale-normalized gene expression values.

### CLDX data assembly, processing and CNA profiling

Gene expression microarray data from the National Cancer Institute MicroXeno project were downloaded from the GEO repository (http://www.ncbi.nlm.nih.gov/geo), under accession number GSE 48433 (ref ^39^). Data were processed as described above. CNA profiles were generated using the FGMP method, and model-acquired CNAs were identified by e-karyotyping, as described above. The *in vitro* cultured (P0) cell line gene expression values were used as reference in the e-karyotyping analysis. The numerical karyotypic complexity categorization of the cell lines was obtained from the publication by Roschke et al. ^61^.

### Cell line data assembly, processing and CNA profiling

Whole-exome sequencing data from 9 newly-derived cell lines were obtained from Tseng et al. (manuscript in preparation). CNA profiles were generated from these data using the ReCapSeg program (http://gatkforums.broadinstitute.org/gatk/categories/recapseg-documentation), from the ratio of tumor read depth to the expected read depth (as determined from a panel of normal samples). Gene expression microarray data were obtained from the GEO repository (http://www.ncbi.nlm.nih.gov/geo). Accession numbers are provided in Supplementary Table 3. Data were processed as described above. CNA profiles were generated using the FGMP method, and model-acquired CNAs were identified by e-karyotyping, as described above. Renal cancer CNA data were obtained directly from the publication by Cifola et al. ^62^, and model-acquired CNAs were identified as described above. For the comparison of model-acquired CNA rates across passages, samples were compared to the earliest available passage (p=0 or p=1). Samples from p<=7 were defined as early passage, samples from p=10 were defined as medium passage, and samples from p>=19 were defined as late passage.

### Drug response association analyses

PDX drug response data were obtained from the publication by Gao et al ^3^. For the analysis of the association between chemotherapy response and absolute levels of aneuploidy, the CNA fraction was determined according to the FGMP-derived CNA profiles of the latest passage sample available from each model. Low CNA levels were determined as CNA fraction<0.3; intermediate CNA levels were determined as 0.3<CNA fraction<0.7; high CNA levels were determined as CNA fraction>0.7. For the analysis of the association between chemotherapy response and the degree of genomic instability (DGI), the DGI level of each model was determined as the number of discrete model-acquired CNAs per passage, using the latest passage sample available from each model: low DGI=0, 0<intermediate DGI<4, high DGI>4. The BestAvgResponse values were used to make response calls. Association tests were conducted in each available tissue type independently, yielding a total of six drug-tissue association tests (representing five chemotherapies in five tissue types). For the analysis of the association between targeted therapy response and the existence of specific arm-level events, the arm-level copy number status of each model was set according to the FGMP-derived CNA profiles of the latest passage sample available from that model. The BestAvgResponse values were used to make response calls. For each of the 12 recurrent TCGA events that tend to disappear throughout PDX passaging, its association with PDX drug response was evaluated in the relevant cancer type. All targeted drugs that were used as single agents, and that showed at least partial response in at least one animal, were evaluated. Drug combinations were also evaluated, if one (or both) of the drugs in the combination was not tested as a single agent. A total of X association tests were performed (representing 15 single agent drugs and five drug combinations in three tissue types).

### Statistical analyses

The significance of the differences in prevalence and rate of absolute CNAs and of model-acquired CNAs between PDX passages, between primary and metastatic PDXs, between P53-WT and P53-mutated/deleted PDXs, between PDXs from the most stable (upper quartile) and least stable (lower quartile) tissue types, between cell line passages, between CLDX passages, and between CLDXs from cell lines of distinct numerical karyotypic complexities, was determined using the two-tailed Wilcoxon rank-sum test. The significance of the difference in similarity scores between “sibling” and “non-sibling”, and the significance of the difference in CNA rates between PDXs and cell lines, were determined using a stratified bootstrap test, permuting the data 100,00 times within each tissue type. The significance of the gene expression signature trends observed throughout PDX passaging was determined by the Kruskal-Wallis rank-sum test. The significance of correlations between PDX and TCGA data was determined using a Spearman’s correlation test. To evaluate the tendency to acquire or to lose recurrent TCGA CNAs during PDX propagation, recurrent CNAs were defined for each tissue type as those that recur in over 40% of the samples, and the number of events that involve these CNAs were computed in the lineage-matched PDX cohorts; the significance of the difference between the emergence frequency and the loss frequency was determined using the McNemar’s test. The significance of the difference in CNA prevalence between early and late passage PDX samples was evaluated using the one-tailed Fisher’s exact test. The significance of the association between chromosome arms and drug response was determined using the Wilcoxon rank-sum test, with FDR multiple test correction performed for each tissue type independently. Box plots show the median, 25^th^ and 75^th^ percentiles, lower whiskers show data within 25^th^ percentile −1.5 times the IQR, upper whiskers show data within 75^th^ percentile +1.5 times the IQR, and circles show the actual data points. Violin plots show the combination of a box plot and a kernel density plot, in which the width is proportional to the relative frequency of the measurements. All of the statistical tests were performed, using the R statistical software (http://www.r-project.org/), and the box plots and violin plots were generated using the ‘boxplot’ and ‘vioplot’ R packages, respectively.

### Code availability

The codes used to generate and/or analyze the data during the current study are publically available, or available from the authors upon request.

### Data availability

The datasets generated during and/or analyzed during the current study are available within the article, its supplementary information files, or available from the authors upon request.

## Legends to Supplementary Figures and Tables

**Supplementary Figure 1: CNA profiles from PDX gene expression data are highly similar to those from PDX SNP array data**

Moving average plots of PDX SNP arrays (upper panels) and their corresponding gene expression arrays (lower panels) in six representative cancer types. The CNAs identified in each sample by our pipeline (**Methods**) are depicted as rectangles above the affected genomic regions. Gains are shown in red, losses in blue.

**Supplementary Figure 2: The CNA landscapes of PDXs are highly similar to those of primary tumors from matched tissues**

CNA frequency plots of PDX model types and the respective primary tumor types from TCGA, showing that PDXs generally exhibit the aneuploidies and CNAs that are characteristic of each tissue type. Gains are shown in red, losses in blue.

**Supplementary Figure 3: Gradual evolution of CNA landscapes throughout PDX passaging**

PDX models acquire CNAs throughout their *in vivo* propagation. **(a)** Bar plots present the fraction of the PDX models with at least one model-acquired CNA, as a function of the number of passages between measurements. (**b**) Box plots present the number of discrete CNAs as a function of the number of passages between measurements. Bar, median; box, 25^th^ and 75^th^ percentiles; whiskers, data within 1.5*IQR of lower or upper quartile; circles: all data points. P-values indicate significance from a Wilcoxon rank-sum test. (**c**) Box plots present the proportion of genes affected by CNAs as a function of the number of passages between measurements. Bar, median; box, 25^th^ and 75^th^ percentiles; whiskers, data within 1.5*IQR of lower or upper quartile; circles: all data points. P-values indicate significance from a Wilcoxon rank-sum test. (**d**) Equal rates of acquiring new CNAs and losing existing ones in PDXs. Violin plots present the absolute CNA fraction of PDX models at early (p<2), medium (2<p<4) and late (p>=4) passages. Bar, median; colored rectangle, 25^th^ and 75^th^ percentiles; width of the violin indicates frequency at that CNA fraction level. n.s., non-significant (Wilcoxon rank-sum test).

**Supplementary Figure 4: PDX models from metastases exhibit larger CNA fractions and higher CIN70 scores than PDX models from primary tumors**

PDX models from metastases are more aneuploid than those from primary tumors. (**a**) Box plots present the absolute CNA fraction of PDX models from primary tumors (n=563) and from metastases (n=98). (**b**) Box plots present chromosomal instability (CIN70) signature scores of PDX models from primary tumors (n=563) and from metastases (n=98). P-values indicate significance from Wilcoxon rank-sum tests.

**Supplementary Figure 5: Expansion of pre-existing subclones during PDX propagation demonstrated by identification of LOH “reversion”**

Alleles that seem to have been lost in early-passage PDX tumors can “re-appear” in later passages of the same PDX models, demonstrating expansion of rare pre-existing subclones throughout PDX propagation. Plots present the loss of heterozygosity (LOH) status along the genomes of four PDX models from Eirew et al. ^7^. LOH events are shown in purple. For each model, shown are two passages. Arrows mark large (>10Mb) chromosomal segments for which LOH was identified at the earlier passage, but both alleles were present at the later passage.

**Supplementary Figure 6: Genomic instability in PDXs correlates both the genomic instability and the heterogeneity levels of primary tumors**

(**a**) The DGI of PDXs and that of primary tumors correlate extremely well. In PDXs, tissue DGI was defined as the median CNA fraction affected per passage. In TCGA tumors, tissue DGI was defined as the fraction of samples with whole-genome duplication (WGD). (**b**) The DGI of PDXs also correlates extremely well with intra-tumor heterogeneity (ITH) of primary tumors (including skin tissue). The DGI of PDXs was defined as the median number of arm-level CNAs per passage. The heterogeneity of primary tumors was defined as the median number of clones per tumor. Spearman’s rho values and p-values indicate the strength and significance of the correlations, respectively.

**Supplementary Figure 7: Disappearance of recurrent CNAs throughout PDX propagation: opposite trends of patient-acquired and model-acquired CNAs**

Twelve recurrent arm-level CNAs, which were observed in >40% of TCGA samples, were found to be preferentially lost during PDX passaging. Heatmaps present the model-acquired arm-level CNAs identified in five PDX tumor types: breast, brain, colon, lung and pancreas. Gains are shown in red, losses in blue. The chromosome arms that show an opposite acquisition trend to that seen in human patients are highlighted with arrows. 86% of these events represent the disappearance of a CNA that existed at an earlier passage, rather than the acquisition of the opposite CNA.

**Supplementary Figure 8: Disappearance of recurrent CNAs throughout PDX propagation: prevalence differences between early and late passages**

Recurrent CNAs that tend to disappear during PDX passaging are less commonly identified in late compared to early passage PDX samples. Absolute CNA frequency plots of three PDX model types (breast, brain and lung) at early and late passage numbers are presented. Gains are shown in red, losses in blue. Nine of the twelve events that tend to disappear in PDXs are less common in high passage PDXs (highlighted by arrows). P-values indicate significance from a Fisher’s exact test.

**Supplementary Figure 9: Genomic instability of PDXs is comparable to that of cell lines and CLDXs**

(**a**) Gradual evolution of CNA landscapes throughout passaging of newly-derived cell lines. Box plots present model-acquired CNA fraction as a function of *in vitro* passage number. (**b**) Similar rates of CNA acquisition in PDXs and in newly-derived cell lines. Dot plots present the distribution of model-acquired CNA fractions across three available cancer types. Cell lines used for this analysis are listed in **Supplementary Table 3**. (**c**) The CNA acquisition rate of CLDXs is associated with the numerical karyotypic complexity of the parental cell lines. Violin plots present the fraction of CNAs acquired by passage 10 as a function of numerical karyotypic complexity. P-values indicate significance from a Wilcoxon rank-sum test.

**Supplementary Figure 10: *De novo* CNAs may play a role in PDX CNA dynamics as well**

(**a**) Model-acquired CNAs keep emerging at high *in vivo* passages. Plots present the model-acquired CNAs in multiple passages of two breast PDX models ^5^. (**b**) Unique events also emerge in “sibling” PDXs, which were derived from the same primary tumor and propagated independently in mice. Plots present the model-acquired CNAs in pairs of passages from breast (PDX2127), lung (PDX1726), pancreas (PDX2081) and skin (PDX1655) PDX models ^3^. Gains are shown in red, losses in blue. (**c**) PDXs with a mutant or deleted p53 present a significantly higher rate of CNA acquisition throughout passaging, compared to their WT counterparts. Box plots present the rate of model-acquired CNAs in PDX models without (PDX-WT; n=65) and with (TP53 mut/del; n=110) a TP53 perturbation. P-value indicates significance from a Wilcoxon rank-sum test.

**Supplementary Table 1: Summary of PDX datasets**

A list of the datasets included in this study, together with their accession numbers, tumor types, the number of PDX models and samples included in them, the experimental platform used, and the Pubmed ID number of the original study that generated them.

**Supplementary Table 2: Comparison of DNA- and RNA-based CNA profiles**

A comparison of model-acquired CNAs inferred from DNA and RNA data from the same tumor samples.

**Supplementary Table 3: Summary of newly-derived cell lines**

A list of the newly-derived cell lines included in this study, together with their accession numbers, tumor types, passage numbers, and the Pubmed ID number of the original study that generated them.

**Supplementary Data 1: CNA profiles of PDX samples**

PDX CNA profiles generated in this study from gene expression data. The first tab provides a full description of the samples. The second tab provides a segmental aberration matrix, in a format readily visualized by the Integrative Genomics Viewer (IGV; https://www.broadinstitute.org/igv/).

**Supplementary Data 2: Model-acquired CNAs in PDX samples**

PDX model-acquired CNAs identified in this study from gene expression data. The first tab provides a full description of the samples. The second tab provides a segmental aberration matrix, in a format readily visualized by the Integrative Genomics Viewer (IGV; https://www.broadinstitute.org/igv/).

**Supplementary Data 3: CNA profiles of CLDX samples**

CLDX CNA profiles generated in this study from gene expression data. The first tab provides a full description of the samples. The second tab provides a segmental aberration matrix, in a format readily visualized by the Integrative Genomics Viewer (IGV; https://www.broadinstitute.org/igv/).

**Supplementary Data 4: Model-acquired CNAs in CLDX samples**

CLDX model-acquired CNAs identified in this study from gene expression data. The first tab provides a full description of the samples. The second tab provides a segmental aberration matrix, in a format readily visualized by the Integrative Genomics Viewer (IGV; https://www.broadinstitute.org/igv/).

## Contributions

U.B.-D. conceived the project, collected the data and carried out the analyses; G.H. assisted with computational analyses; Y.-Y.T. and J.S.B. provided cell line data. C.O. assisted with the copy number analysis of cell lines. B.W. assisted with figure design and preparation. J.S. assisted with the copy number analysis of TCGA samples. R.B. and T.R.G. directed the project. U.B.-D., R.B. and T.R.G. wrote the manuscript.

## Acknowledgements

The authors thank Lude Franke for assistance with functional genomic mRNA profiling; James McFarland for assistance with statistical analysis; Adam Bass, Keith Ligon, Andrew J. Aguirre and Jochen Lorch for providing the samples for cell line derivation; Andrew Tubelli for assistance with figure preparation; Matthew Meyerson, Andrew J. Cherniack, Alison Taylor and Zuzana Tothova for helpful discussions. U.B.-D. is supported by a Human Frontiers Science Program postdoctoral fellowship.

